# Discovery of functional gene markers of bacteria for monitoring hydrocarbon pollution in the marine environment - a metatranscriptomics approach

**DOI:** 10.1101/857391

**Authors:** Kamila Knapik, Andrea Bagi, Adriana Krolicka, Thierry Baussant

## Abstract

The use of natural marine bacteria as “oil sensors” for the detection of pollution events can be suggested as a novel way of monitoring oil occurrence at sea. Nucleic acid-based devices generically called genosensors are emerging as potentially promising tools for *in situ* detection of specific microbial marker genes suited for that purpose. Functional marker genes are particularly interesting as targets for oil-related genosensing but their identification remains a challenge. Here, seawater samples, collected in tanks with oil addition mimicking a realistic oil spill scenario, were filtered and archived by the Environmental Sample Processor (ESP), a fully robotized genosensor, and the samples were then used for post-retrieval metatranscriptomic analysis. After extraction, RNA from ESP-archived samples at start, day 4 and day 7 of the experiment was used for sequencing. Metatranscriptomics revealed that several KEGG pathways were significantly enriched in samples exposed to oil. However, these pathways were highly expressed also in the non-oil-exposed water samples, most likely as a result of the release of natural organic matter from decaying phytoplankton. Temporary peaks of aliphatic alcohol and aldehyde dehydrogenases and monoaromatic ring-degrading enzymes (e.g. *ben*, *box*, and *dmp* clusters) were observed on day 4 in both control and oil tanks. Few alkane 1-monooxygenase genes were upregulated on oil, mostly transcribed by families *Porticoccaceae* and *Rhodobacteraceae*, together with aromatic ring-hydroxylating dioxygenases, mostly transcribed by *Rhodobacteraceae*. Few transcripts from obligate hydrocarbonoclastic genera of *Alcanivorax*, *Oleispira* and *Cycloclasticus*, were significantly enriched in the oil-treated tank in comparison to control, and these were mostly transporters and genes involved in nitrogen and phosphorous acquisition. This study highlights the importance of seasonality, *i.e.*, phytoplankton occurrence and senescence leading to organic compound release which can be used preferentially by bacteria over oil compounds, delaying the latter process. As a result, such seasonal effect can reduce the sensitivity of genosensing tools employing bacterial functional genes to sense oil. A better understanding of the use of natural organic matter by bacteria involved in oil-biodegradation is needed to develop an array of functional markers enabling the rapid and specific *in situ* detection of anthropogenic pollution.

## 1 Introduction

The use of ecogenomic sensors and devices that can automatize molecular analytical techniques are promising for *in situ* marine applications, such as understanding fundamental processes in complex and dynamic environments, detection of episodic events such as oil leakages from subsea operational infrastructures and monitoring of ecosystem recovery (Ussler, Preston et al. 2013, Danovaro, Carugati et al. 2016, Ottesen 2016, Jones, Gates et al. 2019). The development of genosensors that specifically target the genetic material of microorganisms that respond to oil for use as oil sensors and their automatization for *in situ* monitoring of pollution events at sea is novel (Palchetti, Bettazzi et al. 2018, Bagi, Knapik et al. 2019).

Microbial communities inhabiting the water column of the global ocean are appropriate for the development of genosensors as these communities are ubiquitous and respond rapidly to changes in their environment (Nogales, Lanfranconi et al. 2011). In a recent study, bacteria were found to be the most responsive to drilling and production operations in the South Taranaki Bight region of the Tasman Sea, further emphasizing the need for genosensing targets based on this domain of life (Laroche, Wood et al. 2018). Next-generation sequencing (NGS)-based approaches targeting microbes, or more specifically bacteria in aquatic environments, have been suggested and explored as the new frontier in water quality assessment (Tan, Ng et al. 2015). The assessment and evaluation of this type of novel approach towards integration into routine monitoring practices has been increasing in the past decade (Hering, Borja et al. 2018, Cordier, Lanzén et al. 2019). NGS-based analysis of microbial response can also be used to identify potential bioindicator species and genes. Quantitative molecular assays targeting these can then be developed and integrated into ecogenomic devices.

Ecogenomic devices equipped with molecular assays targeting specific biomarker genes, could have a significant impact for application in industrialized areas of the ocean where legal requirements governing anthropogenic activities call for novel technologies with quick response times to mitigate their effect on the ocean ecosystem.

One off-the-shelf genosensing device is the Environmental Sampling Processor (ESP), a fully developed and automated ecogenomic sensor capable of performing different analytical assays at sea for remote detection of marine microbes, microalgae and small invertebrates. This unique device is the product of many years of development during which the ESP has evolved from a so-called 2G-ESP (large, fixed on a buoy or a drifter) to the present 3G-ESP (smaller, mobile in a LR-AUV) (Pargett, Birch et al. 2015, Scholin 2017). While the ESP device cannot perform *in situ* RNA sequencing, metatranscriptomic analysis of ESP archived samples have been performed to uncover *in situ* patterns of microbial activity (Ottesen, Marin et al. 2011). Although the ESP was not developed for the purpose of environmental monitoring of offshore industrial activities such as Oil & Gas, two recent studies indicated it could potentially be used in that context by quantifying taxonomic markers of oil pollution using specific 16S rRNA genes (Krolicka, Boccadoro et al. 2014, Krolicka, Boccadoro et al. 2019).

Oil contamination is known to induce characteristic shifts in the composition of pelagic marine bacterial communities resulting in higher abundance of some species within the Gamma- and Alphaproteobacteria classes (Harayama, Kasai et al. 2004, Chakraborty, Borglin et al., McGenity, Folwell et al. 2012, Kostka, Teske et al. 2014, Brakstad, Throne-Holst et al. 2015). There are several bacterial groups which have been suggested as indicators of oil pollution at sea. Typical microbial species associated with marine oil spills in different environments are alkane-degrading microbes such as *Alcanivorax*, *Oleispira* and *Colwellia*, which are succeeded by PAH-degrading groups such as *Cycloclasticus* and *Marinobacter* (Hazen, Dubinsky et al. 2010, Dubinsky, Conrad et al. 2013, Brakstad, Throne-Holst et al. 2015, Ribicic, Netzer et al. 2018). Among them *Oleispira* and uncultured *Oceanospirillaceae* were identified as potentially good indicators for oil pollution in cold waters (Krolicka, Boccadoro et al. 2014, Gontikaki, Potts et al. 2018) responding fast and increasingly to oil load in water. These 16S rRNA gene-based amplicon sequencing studies are used to characterize the composition of these communities and to follow their temporal succession. In comparison to metagenomics or metatranscriptomics, the 16S rRNA-based approach is less sensitive to pitfalls on the bioinformatic analysis side and thus allows for a more reliable comparative analysis between samples. However, the full potential of the phylogenetic marker-based analysis for oil detection can only be explored when a significant increase in obligate hydrocarbonoclastic bacteria (OHCB) is observed (Yakimov, Timmis et al. 2007). This is because OHCB can utilize only a very narrow range of carbon substrates (*i.e.* hydrocarbons), and hence their bloom can directly be linked to the occurrence of oil in water. Besides oil-degraders, there are hydrocarbon-tolerant opportunistic bacteria which can also benefit from the occurrence of oil in seawater, even though they are not directly involved in oil biodegradation. The drawback of the phylogenetic marker-based analysis is the lack of a direct link to active metabolic pathways which provide a more precise picture about ongoing processes. Therefore, functional markers of such specific pathways related to petroleum degradation are better suited to track occurrence of oil in water directly and can be explored for oil sensing (Zhang, Hu et al. 2018). Some studies have used quantitative functional gene-based biomarker approaches such as qPCR to follow bioremediation processes (mostly soil and sediment) using primers designed for specific hydrocarbon degradation-relevant genes, such as alkane 1-monooxygenase and ring-hydroxylating dioxygenases (Cébron, Norini et al. 2008, Scoma, Hernandez-Sanabria et al. 2015, Shahsavari, Aburto-Medina et al. 2016). Other functional gene marker-based environmental assessments have been mainly limited to DNA-based GeoChip-microarray analysis or other microarray systems (Lu, Deng et al. 2012, Vilchez-Vargas, Geffers et al. 2013). The potential of using mRNA transcripts of genes involved in oil biodegradation as biomarkers of oil pollution has not been fully explored (Bargiela, Yakimov et al. 2017). Only a few studies have attempted to pursue a metatranscriptomic approach in relation to marine hydrocarbon pollution (Mason, Hazen et al. 2012, Rivers, Sharma et al. 2013, Yergeau, Maynard et al. 2015, Vikram, Lipus et al. 2016, Tremblay, Yergeau et al. 2017, Tremblay, Fortin et al. 2019).

In the present study, two 2G-ESP instruments were used in a laboratory-controlled setup to analyze and preserve bacterioplankton samples from seawater collected during an oil exposure experiment mimicking a realistic oil spill condition. The samples were used to perform post-retrieval metatranscriptomic analyses for characterizing the expressed genes of marine microbial communities in response to oil. The main aim of this study was to identify a set of key genes differentiating control from oil treated seawater samples, with the vision to develop specific functional gene-based assays compatible with ESP, for real-time *in situ* detection. The metatranscriptomics analysis was carried out to perform: (1) a taxonomic investigation of the transcripts to identify the pool of active bacteria increasing significantly in oil treated seawater and (2) a differential expression analysis of genes between control and oil exposed water samples to identify a pool of functional genes directly or indirectly involved in oil compound biodegradation. The main hypotheses tested in this study were that in the presence of oil (1) taxonomic and functional profile of seawater becomes distinct from that of control, (2) there is an increase in expression of hydrocarbon specific initial oxygenase genes involved in the activation of aliphatic and aromatic hydrocarbons, and (3) there is an increase in the abundance of genes from OHCB. The outcome of this study is a milestone to support further development toward the use of novel ecogenomic sensors such as ESP for monitoring transient oil pollution events from offshore industrial activities.

## 2 Materials and Methods

### 2.1 Experimental setup

Seawater was collected though a pipeline from approximately 20 m depth (temperature and salinity was approximately 8°C and 33 PSU, respectively) from a pristine coastal area near Kvitsøy (59°03′44″N 05°24′42″E) (Byfjorden, Norway) in a 1 m^3^ container in April 2016 (spring season). The sample was transported to the NORCE facility (Randaberg, Norway) within approximately 2 hrs. Clean 150 L tanks were filled with the Kvitsøy seawater and allowed to acclimate in the laboratory for 24 h at 5°C in the dark prior to the start of the experiment. The experimental exposure system was based on the setup described in (Arnberg, Calosi et al. 2018). Oil treated tanks (O) were prepared in triplicate by adding a pre-mixed stock solution of oil (a naphthalenic crude oil from the Troll oil field) and seawater to the tanks to obtain a nominal oil concentration of 10 mg·L^-1^ at the start of the experiment. At the center of each tank, a funnel attached to a water pump (Aquaclear 70, Hagen, USA) was used to pull water and oil downwards in the tank and allow for sufficient energy to mix water and oil. There was no lid on the tanks to allow for natural weathering from evaporation of the most volatile compounds as this would occur in the field. Control tanks (C) were subsequently prepared with the same funnel/pump system without addition of oil and were placed in a separate climate room also set at 5 °C to avoid contamination from the evaporated oil compounds. For each control and oil exposed treatment, three tanks were used but only one for each treatment was dedicated to the present ESP metatranscriptomic study. The experiment was conducted in darkness.

### 2.2 Sampling

#### 2.2.1 Bacterioplankton analysis

Seawater samples were collected manually in autoclaved glass bottles from the control and oil treated tanks daily for 7 days. One sample of 1000 mL of water was collected using a pipe placed in the middle of each tank at the start of the experiment (T0, after 24 hours of acclimation) and then 700 mL was collected each following day for 7 days. Seawater samples were immediately brought to two ESPs. The glass bottles were connected to the ESP instrument which processed the samples automatically. The ESPs filtered the samples onto 25 mm diameter, 0.22 µm pore size hydrophilic Durapore filters (GV, Millipore) and archived the filters after addition of RNA*later*™ stabilization solution (Ambion) for RNA preservation (Ottesen, Marin et al. 2011) (Preston, Harris et al. 2011). After each operation, the filters were retrieved from the ESP and stored at -80°C until use. In total, 15 filters were collected during the experiment (1 filter/day/condition and the initial seawater) and 5 were selected for the metatranscriptomics analysis presented here.

#### 2.2.2 Chemical analysis

Seawater was collected from each of the tanks for total hydrocarbon content (THC) and polycyclic aromatic hydrocarbon (PAH) composition by GC-FID and GC-MS analysis, respectively, on day 1, 3, 5 and 7. PAH analysis was performed for the 16 EPA (Environmental Protection Agency, USA) priority aromatic compounds (Keith 2015). All analyses were performed by Intertek West Lab AS (Tananger, Norway) according to ISO 28540:2011. The limit of quantification for THC analysis was 0.4 mg·L^-1^ and for PAHs, this ranged from 0.01 to 0.02 µg·L^-1^.

### 2.3 Nucleic acid extractions and sequencing

Total RNA and DNA was extracted from five archived filters for metatranscriptomics and amplicon sequencing analysis collected after 24 hr acclimation (T0), 4 (T4C, T4O) and 7 days (T7C, T7O). The extraction was carried out using the AllPrep Bacterial DNA/RNA/Protein Mini Kit (Qiagen) following the manufacturer’s instructions. RNA concentration was quantified on a Qubit instrument (Thermo Fisher Scientific) and the quality of the samples was evaluated on agarose gel.

#### 2.3.1 Metatranscriptomics

Total RNA (2-8 µg) (details in Table S1) was sent to the sequencing facility at the University of Cambridge (Cambridge, UK) where it was subjected to Ribo-Zero ribosomal RNA removal prior to library preparation using Illumina TruSeq Stranded mRNA kit (Illumina). Paired-end (75 bp) sequencing was carried out on a NextSeq500 platform using the 150-cycle kit.

#### 2.3.2 Amplicon sequencing

The SSU rRNA gene was amplified using forward Bakt_341F (5’-CCTACGGGNGGCWGCAG-3’) and reverse primer Bakt_805R (5’-GACTACHVGGGTATCTAATCC-3’) primers, which target the V3 and V4 region (Mizrahi-Man, Davenport et al. 2013, Sinclair, Osman et al. 2015). The PCR master mix consisted of 12.5 μL of AmpliTaq Gold® 360 PCR Master Mix (Thermo Fisher Scientific, Carlsbad, California, USA), 1 μL of each primer to the final concentration of 250 nM, 2.5 μL of GC enhancer (Thermo Fisher Scientific, Carlsbad, California, USA), 2 μL of DNA (1:10 diluted) and double-distilled water (ddH_2_O) to the final volume of 25 μL. The reaction cycling conditions were: 95°C for 10 min, followed by 31 cycles of 95°C for 30 s, 54°C for 45 s, 72°C for 7 min, with a final extension step at 72°C for 7 min. To ensure amplification of uncontaminated products, all PCRs included negative controls (no template samples). The amplicons were prepared in triplicates per individual sample and then merged, purified using PureLink™ Quick Gel Extraction and PCR Purification Combo Kit (Thermo Fisher Scientific, Carlsbad, California, USA) and finally quantified using a Qubit® Fluorometer (Thermo Fisher Scientific, Carlsbad, California, USA). The prepared amplicons were sequenced from both ends at BaseClear BV (Netherlands) using the Illumina MiSeq platform. The raw sequencing reads are publicly available in the Sequence Read Archive (SRA) NCBI within the PRJEB32487 study under the accession numbers ERS3402758-ERS3402761.

### 2.4 Data analysis

#### 2.4.1 Metatranscriptomics

The entire bioinformatic workflow is presented in Figure S1. Raw reads were processed as single reads. Quality filtering and trimming was done using PRINSEQ and Trimmomatic. PRINSEQ removed low quality reads (PHRED < 10), duplicates, reads with ambiguous bases (N) and reads with low complexity (Entropy < 70) (Schmieder and Edwards 2011). Trimmomatic (Bolger, Lohse et al. 2014) was used to trim (sliding window of 4:15) and remove reads shorter than 50 bp. The next step of the read preprocessing was ribosomal RNA (16S/18S, 23S/28S and 5S rRNA) removal with the *SortMeRNA tool (Kopylova, Noe et al. 2012) using the SILVA (Burge, Daub et al. 2013) and Rfam (Quast, Pruesse et al. 2013) databases*.

High-quality non-ribosomal RNA reads from all five samples were then combined in one file and *de novo* assembled with IDBA-Tran using default parameters (Peng, Leung et al. 2013). Only those reads which mapped back to these contigs were used in downstream analysis in order to decrease the probability of spurious annotations of short reads (Toseland, Moxon et al. 2014). Contigs were used to assess the taxonomic composition of the metatranscriptome. Protein coding DNA sequences (CDS, i.e., genes – these terms will be used interchangeably in the text) were predicted from contigs using MetaGeneMark (Zhu, Lomsadze et al. 2010). The read counts of assembled transcripts and putative CDS were determined by mapping the quality-filtered reads of each sample against the assembled contigs and CDS using BWA (Li and Durbin 2009). Reads which did not map back to contigs or CDS were not used in further analysis. The read counts of transcripts and CDS sequences were normalized as transcripts per million (TPM) according to Wagner et al. (Wagner, Kin et al. 2012).

The statistics of each read processing step are shown in Table S2. The raw sequencing reads are publicly available in the Sequence Read Archive (SRA) NCBI under the project number PRJNA556155.

For taxonomic classification, contigs were aligned to the NCBI non-redundant (nr) protein database (ftp.ncbi.nlm.nih.gov/blast/db/FASTA/nr.gz) using DIAMOND v0.9.9.110 (Buchfink, Xie et al. 2015) and blastx with default parameters. The blastx results were analyzed using the longReads LCA algorithm of MEGAN6 with parameters of percent to cover = 80.0, min score = 50.0, max expected = 0.001, top percent = 10.0, min support percent = 0.0, and min support = 1 (Huson, Auch et al. 2007). MEGAN assignments were then exported using the ‘taxonName to readName’ separator to obtain a list of contig IDs that aligned to each taxon. In order to obtain the abundance for each taxon in each sample, contig IDs (‘readName’) were replaced by TPM counts using a custom script. STAMP was used to generate a principal component analysis (PCA) plot based on relative abundance of families with sum relative abundance > 0.001% (Parks, Tyson et al. 2014).

Functional annotations of all CDS were generated with DIAMOND against the NCBI nr database and using the online KEGG GhostKOALA annotation pipeline (Kanehisa, Sato et al. 2016). STAMP was used to generate a PCA plot based on the TPM counts of unique KEGG orthologs (6,548 KOs).

To identify oil-specific KOs, Venn diagram analysis was performed using the R package VennDiagram (https://CRAN.R-project.org/package=VennDiagram) (Chen and Boutros 2011). Briefly, a list of KOs with TPM > 0 was generated for each sample, then unique and shared KOs were identified and retrieved.

In order to identify CDS that were significantly more abundant in oil treated samples, differential expression analysis was carried out. The raw read count table for CDS was used as input for the DESeq2 available in R/Bioconductor package (Love, Huber et al. 2014). As differential expression analysis requires a minimum of 2 replicates for each condition, the day 4 and day 7 datasets for each condition (control and oil exposed) were treated as replicates, *i.e.*, oil samples (T4O and T7O) were compared with no-oil samples (T4C and T7C). CDS with log2 fold change (log2 FC) values of ≥ 1 and p-values of < 0.05 were defined as significantly differentially abundant/expressed. The putative function of upregulated CDS was determined using the online KEGG GhostKOALA annotation pipeline (Kanehisa, Sato et al. 2016). The result of this analysis was a table of genes, annotated with KEGG Orthology (KO) terms organized according to the KEGG PATHWAY Database (https://www.genome.jp/kegg/pathway.html). KO terms that were not in the KEGG pathways but were relevant to oil degradation, Bacterial motility proteins (ko02035), Secretion system (ko02044) and Transporters (ko02000), were manually collected according to the KEGG BRITE Database (https://www.genome.jp/kegg/brite.html). A custom script was used to obtain the TPM count table, based on the list of CDS IDs corresponding to each KO term. Upregulated CDS with functions linked to hydrocarbon degradation were taxonomically annotated using blastp search with DIAMOND using the combination of the best blast hit and the LCA algorithm (--top parameter set to 2%).

#### 2.4.2 Amplicon sequencing

The raw sequences were processed using USEARCH (v10.0.240_i86linux32) according to the recommended settings (Edgar 2010, Edgar, Haas et al. 2011). The sequences were merged and filtered according to length and quality criteria and sequencing reads that did not meet the quality requirements were discarded. Unique sequences were extracted, then OTU clustering and chimera identification steps were performed. Singletons were removed and finally, reads were mapped to the final OTUs list to assign abundances to each OTU. The OTU table was then rarefied to a depth of 20,000 reads per sample. Unique bacterial OTUs we classified using The Ribosomal Database Project (RDP) Naive Bayesian (Wang, Garrity et al. 2007) database (Edgar R., rdp_16s_v16.fa) and OTUs taxonomy was merged with the rarefied OTU table.

## 3 Results

### 3.1 Chemistry

THC was below detection limit in the acclimated seawater (T0) and in all the control seawater samples. Hydrocarbon analyses confirmed the presence of petroleum hydrocarbons in the seawater exposed to oil on day 1 (1.9 mg**·**L^-1^ THC, 6.04 µg**·**L^-1^ sum of 16 EPA PAHs, and 54.79 µg**·**L^-1^ sum naphthalenes, phenanthrenes, and dibenzothiophenes (NPDs)). Thereafter, the concentration of hydrocarbons appeared to decrease rapidly over time with the largest drop in concentrations taking place between day 1 and 3 (Figure 1, Table S3). As much as 88 and 92% of sum PAHs and sum NPDs disappeared at day 7, respectively, while THC decreased by 73%.

**Figure 1.**
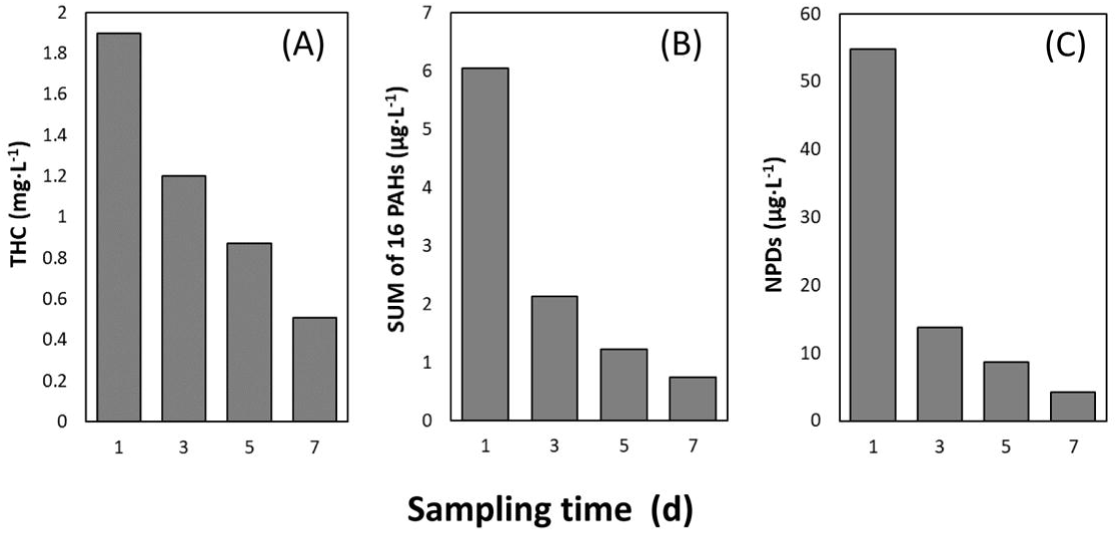
Chemical analysis results. (A): concentrations of total hydrocarbon compounds (THC, mg·L-1), (B): the sum of 16 EPA priority polycyclic aromatic hydrocarbons (SUM of 16 PAHs, µg·L-1) and (C): the sum of naphthalenes, phenanthrenes and dibenzothiophenes (NPDs, µg·L-1) determined by gas chromatography in the oil exposed tank over time.

### 3.2 Sequencing statistics

#### 3.2.1 RNA-Sequencing

Illumina sequencing of the total mRNA from the 5 samples resulted in a total of 127 million quality-filtered reads with ∼17-29 million reads passing QC and rRNA filtering per sample (Table S2). Co-assembly of metatranscriptomic reads produced 628,164 contigs with sizes ranging from 200 bp to 33,000 bp and with an N50 of 521 bp. On average, 69% (67.7-71.1%) of reads did map back onto these contigs. The only exception was the acclimated seawater (T0), for which this proportion was much lower (34.7%).

#### 3.2.2 Amplicon sequencing

The SSU rRNA amplicon sequencing resulted in 37,064-42,712 high quality reads per sample. Both low quality sequencing reads (< 5%) and chimeric sequences (< 0.1%) were removed. In total, 200 OTUs belonging to the domain Bacteria were identified and classified using RDP Classifier.

Rarefaction was carried out to a sequencing depth of 20,000 reads and the resulting OTU table was normalized by converting read counts into relative abundances.

### 3.3 Taxonomic overview

According to the taxonomic analysis of contigs from the metatranscriptome, the seawater communities were primarily dominated by bacteria, except for the acclimated seawater sample (T0) which had a high relative abundance of eukaryotic (∼25%) and unclassified (∼32%) sequences (Figure 2). The fraction of unclassified sequences was below 11% for all other samples.

**Figure 2.**
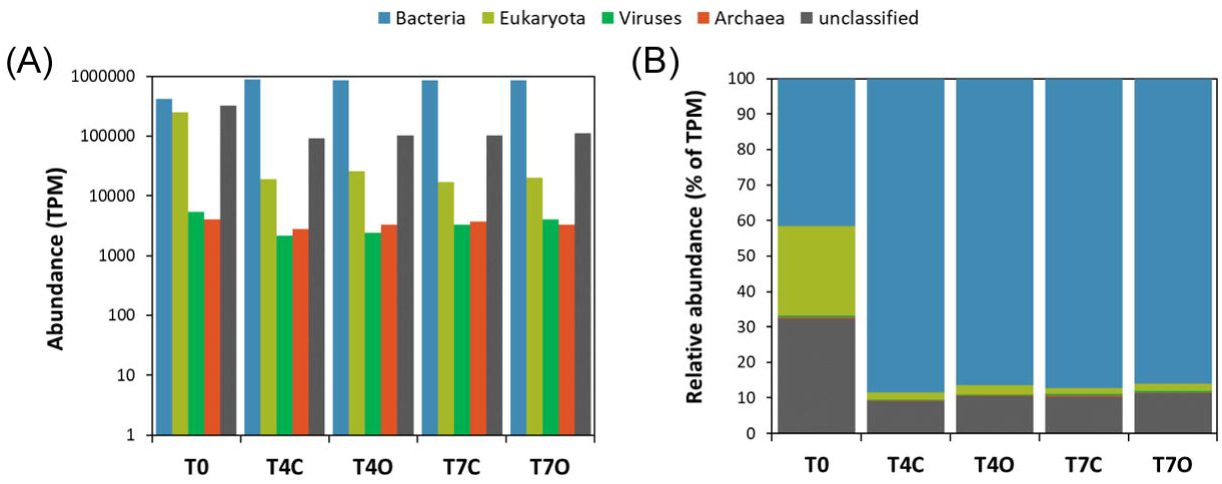
Domain level composition of assembled contigs. TPM counts (A) and relative abundances (B).

The highest diversity on the family level was found in the acclimated seawater (T0) sample, while all other samples had lower Shannon indices (Table S4). When comparing control and oil exposed seawater communities, Shannon indices for the controls were slightly higher than for the oil treatments based on the amplicon sequencing data.

Principal component analysis (PCA) of the relative abundance of families suggested a clear separation of the acclimated seawater from all the other samples along PC1 (81.1% of the variation) while control and oil exposed samples separated along PC2 (17.8% of variation) (Figure 3). The temporal aspect of the differences between the two control and the two oil exposed samples seemed to be less pronounced (PC3, 0.6% of variation).

**Figure 3.**
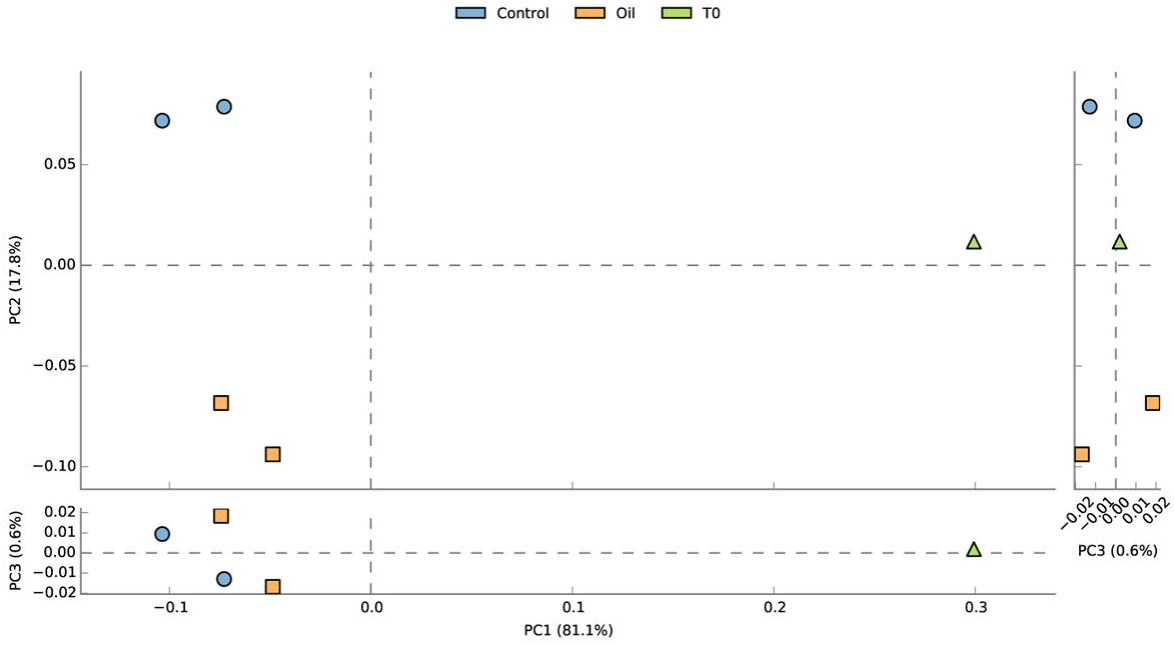
Principal component analysis (PCA) on the family level composition of assembled contigs generated using STAMP. Color codes: blue circles = oil exposed, yellow squares = control, and green triangle = acclimated seawater.

The taxonomic composition of the seawater bacterial communities based on assembled contigs was overall very similar to that based on 16S amplicon sequences (Figure 4, Table S5). The acclimated seawater (T0) was dominated by families *Rhodobacteraceae* (24.1%), *Flavobacteriaceae* (17.8%), other unclassified Bacteria (16.4%), unclassified Gammaproteobacteria (9.3%), OMG group (7.5%), and unclassified Bacteroidetes (5.3%). Compared to the T0 sample, major changes at the family level were revealed by the enrichment of seawater with *Colwelliaceae* (mainly *Colwellia*), *Alteromonadaceae*, and *Campylobacteraceae* (mainly *Arcobacter)* related sequences (Figure 4).

**Figure 4.**
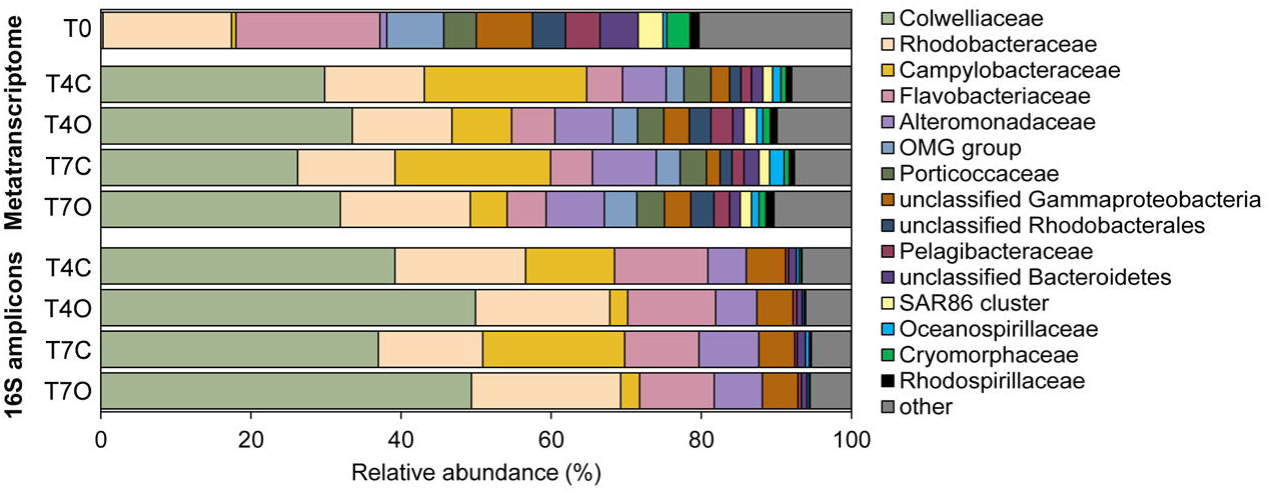
Microbial community composition in seawater exposed to oil and seawater control observed at the start of exposure (T0), at day 4 and day 7 of the exposure determined by metatranscriptomic and 16S rRNA amplicon sequencing analysis. The community composition identified based on the metatranscriptome was derived from the family level taxonomy of annotated contigs (proportion (%) of TPM).

All control and oil-exposed samples collected on day 4 and 7 were dominated by *Colwelliaceae* (23.3-30.7%), *Campylobacteraceae* (3.1-14.9%), *Alteromonadaceae* (9.8-13.7%), *Rhodobacteraceae* (15.8-19.6%), and *Flavobacteriaceae* (4.7-5.4%) (Figure 5). Families *Colwelliaceae*, *Campylobacteraceae* and *Alteromonadaceae* were nearly undetectable in the acclimated seawater (T0 sample) but appeared to bloom in both control and oil exposed incubations. Differences between oil-treated and control samples were rather small with only one of the dominant families, *Campylobacteraceae* (mainly the genus *Arcobacter*) showing a rather clear preference for unexposed seawater (difference in means of control vs oil: 14.8%, excluding T0). *Flavobacteriaceae* was slightly more abundant in T4O sample, while *Alteromonadaceae*, *Rhodobacteraceae*, and *Colwelliaceae* had higher relative abundance in both T4O and T7O in comparison to controls. In addition to the most dominant families, unclassified Gammaproteobacteria, OMG group, unclassified Rhodobacterales, *Pelagibacteraceae*, *Rhodospirillaceae, Cryomorphaceae* and other unclassified Bacteria also had slightly higher relative abundance in oil treated seawater at both timepoints. Unclassified Bacteroidetes and *Oceanospirillaceae* were slightly more abundant in control seawater.

**Figure 5.**
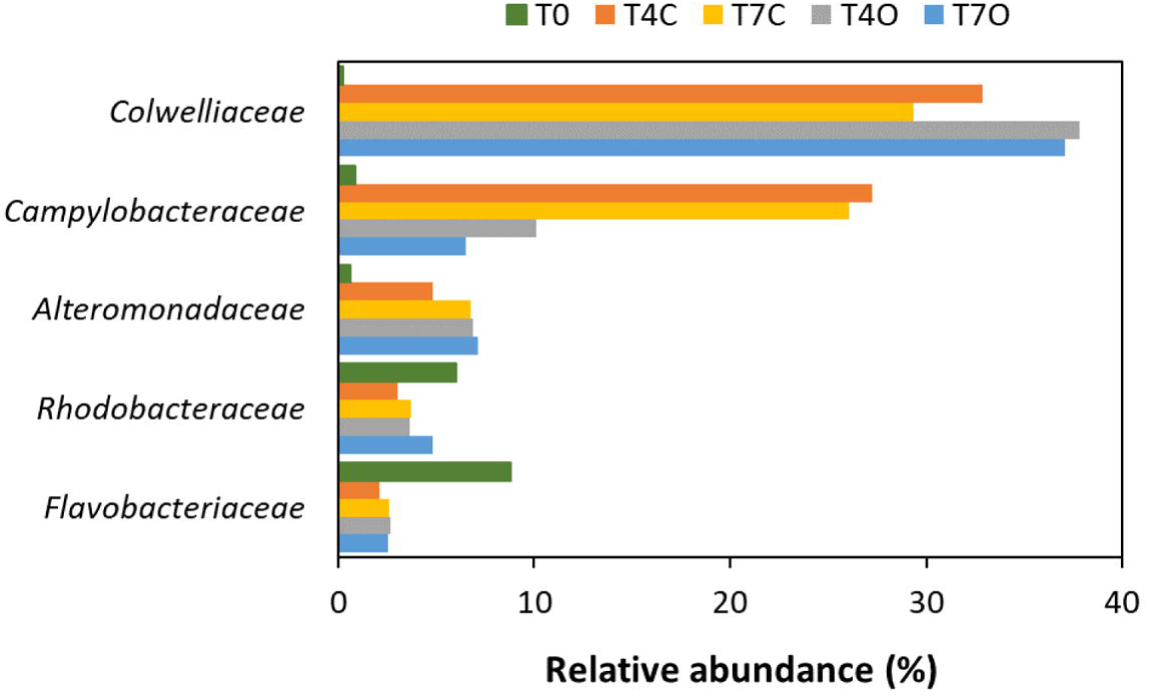
Temporal variation of the relative abundances of the five dominant families.

### 3.4 Functional overview

MetaGeneMark detected 674,434 putative coding sequences (CDS or genes), of which 568,144 were successfully annotated against the NCBI protein database (308,553 unique NCBI IDs) and 176,757 against the KEGG Ortholog Database (6,548 unique KEGG orthologs) (Table S6). PCA of the unique KEGG orthologs revealed similar patterns as observed from the taxonomical analysis, with PC3 (representing temporal separation between samples) explaining 6.4% of the variation (Figure 6). Functional differences between the day 4 and the day 7 oil exposed samples seemed to be more pronounced than taxonomical differences, according to their larger distance along PC3.

**Figure 6.**
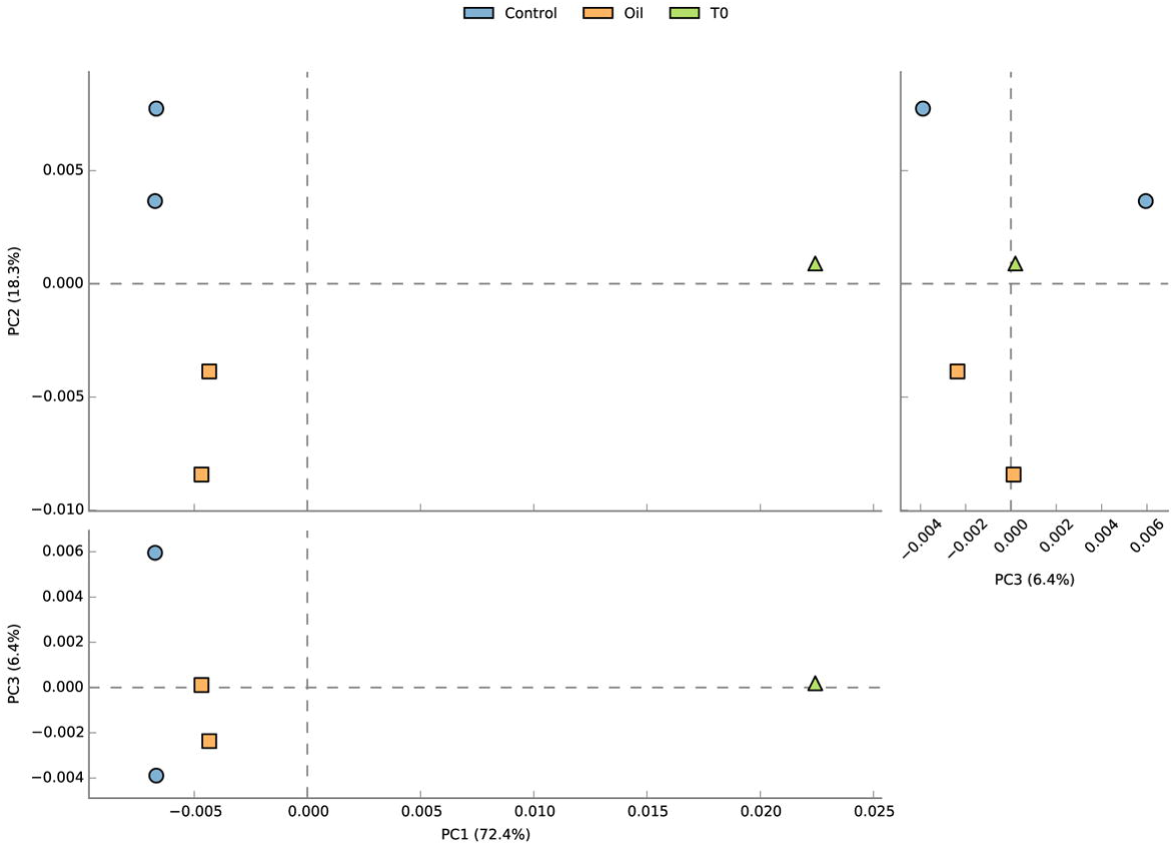
Principal component analysis (PCA) on KEGG ortholog composition, generated using STAMP. Color codes: blue circles = oil exposed, yellow squares = control tank, and green triangle = acclimated seawater.

Based on the most abundant functions, control samples (T4C and T7C) were distinguished from oil exposed ones (T4O and T7O) and the T0 sample by a number of phosphorous acquisition related genes (alkaline phosphatase, phosphate-binding protein, phosphate ABC transporter, phosphonate ABC transporter), carbon and energy storage related functions (phasin family protein and isocitrate lyse) and by the higher abundance of cation acetate symporter genes expressed mainly by *Arcobacter*, *Glaciecola, Sulfitobacter* and *Colwellia* species. The later (day 7) control sample was particularly enriched in these types of genes. Abundant functions differentiating the oil exposed samples (T4O and T7O) from the rest were mainly PQQ-dependent dehydrogenases (methanol/ethanol family) and aldehyde dehydrogenases from different *Colwellia* species. In addition, a TonB−dependent receptor from *Colwellia polaris* and a porin from Candidatus *Colwellia aromaticivorans* was also highly enriched in oil-exposed samples. These transcripts were also present in the control samples, with TPM values 1-13 times lower than in the oil exposed samples. TPM levels in T0 for these transcripts was however very low (TPM < 2). Most of the abundant functions observed in the oil exposed samples showed decreasing TPM values between day 4 and day 7, except for a number of hypothetical proteins and phage associated sequences which increased substantially in the oil exposed tank by day 7.

### 3.5 Upregulated functions

Differential expression analysis (log2 fold-change ≥ 1, *p* ≤ 0.05) resulted in 13,427 CDS (∼2% of all the predicted CDS) that were significantly upregulated in seawater with oil in comparison to control (Table S7). Using the KEGG database, 7,067 differentially expressed (DE) genes (52.6%) were successfully annotated to 1,581 unique KOs, while the BLAST search against the NCBI protein database annotated 11,950 DE genes (89%) to 9,057 unique NCBI IDs.

KEGG mapping of the DE genes revealed that several pathways potentially involved in response to petroleum hydrocarbons were significantly enriched (p < 0.05), including ‘Xenobiotics biodegradation and metabolism’, ‘Fatty acid degradation’ and ‘Degradation of aromatic compounds’ (Figure 7, Table S8). Analysis of the genes within the ‘Xenobiotics biodegradation and metabolism’ (283 genes) and ‘Degradation of aromatic compounds’ (84 genes) revealed gene clusters for catabolism of benzoate (*ben*, *box* and *pca* genes), benzene (*dmp*), catechol (*dmp*, *bph*, *cat*) and homoprotocatechuate (*hpa*). Within the pathway ‘Fatty acid degradation’ (170 genes), gene clusters involved in fatty acid beta-oxidation (*fad*) were expressed (Figure 7). A total of 1,619 genes (12.1% of DE genes) were annotated by KEGG as transporter proteins. Among this large number of transporters, we found those involved in mineral and organic ion transport, ion channels, especially those involved in ammonium transport, as well as various amino acid, phosphate, and sugar transporters. Bacterial motility genes were represented by as many as 219 genes (1.6% of DE genes). The majority of these genes coded for proteins involved in flagella and pilus assembly. Secretion system (107 genes, 0.8%) showed that proteins were secreted mainly through the Sec system, type I and II. Genes associated with two-component system (100 genes, 0.7%) were involved in nitrogen regulation, polar flagellar synthesis, central carbon metabolism, phosphate starvation response, swarming activity and biofilm formation. Biofilm formation genes (87 genes, 0.6%) were associated with transcriptional regulators such as *rpoS* and *oxyR*. However, as Figure 7 shows, these functions were also expressed in the control tank and with the exception of transporters (ko02000), all of them increased in both the control and oil exposed tank in comparison to the T0 sample. Moreover, expression levels of these DE functions largely followed a similar pattern of peaking on day 4 and declining by day 7 both in control and oil exposed tank.

**Figure 7.**
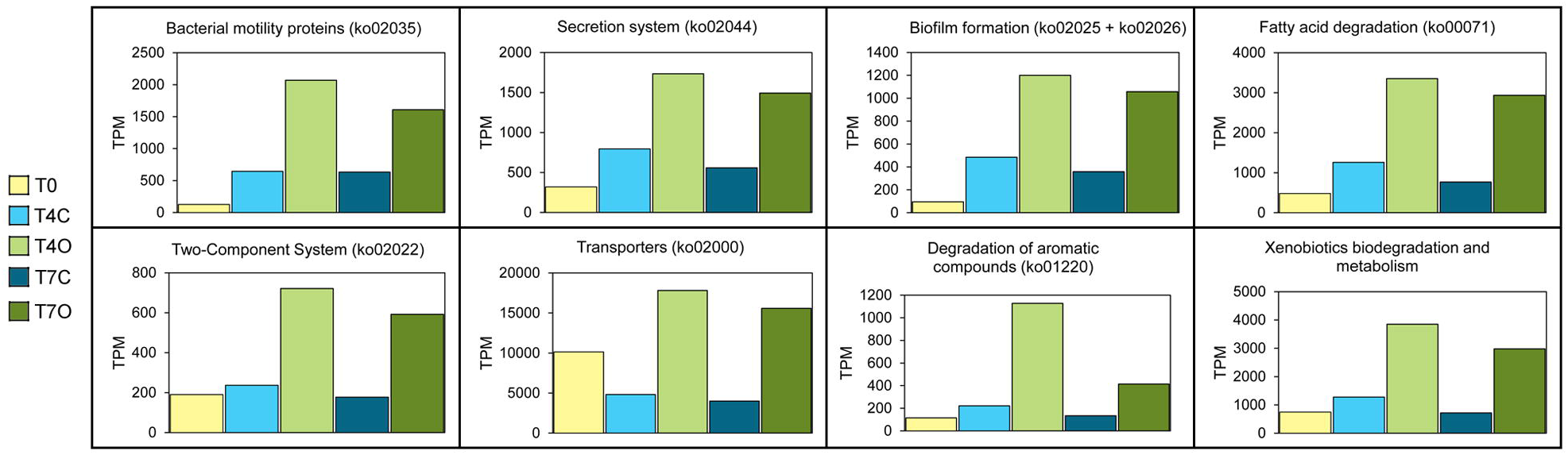
TPM abundances of selected KEGG pathways and BRITE hierarchies of upregulated genes (log2 fold-change ≥ 1, p ≤ 0.05) between control and oil exposed samples.

In order to investigate whether taxonomic profiles could uncover more oil-specific responses, proteins based on KEGG identifiers and genes identified in literature were selected and further investigated (Figure 8).

**Figure 8.**
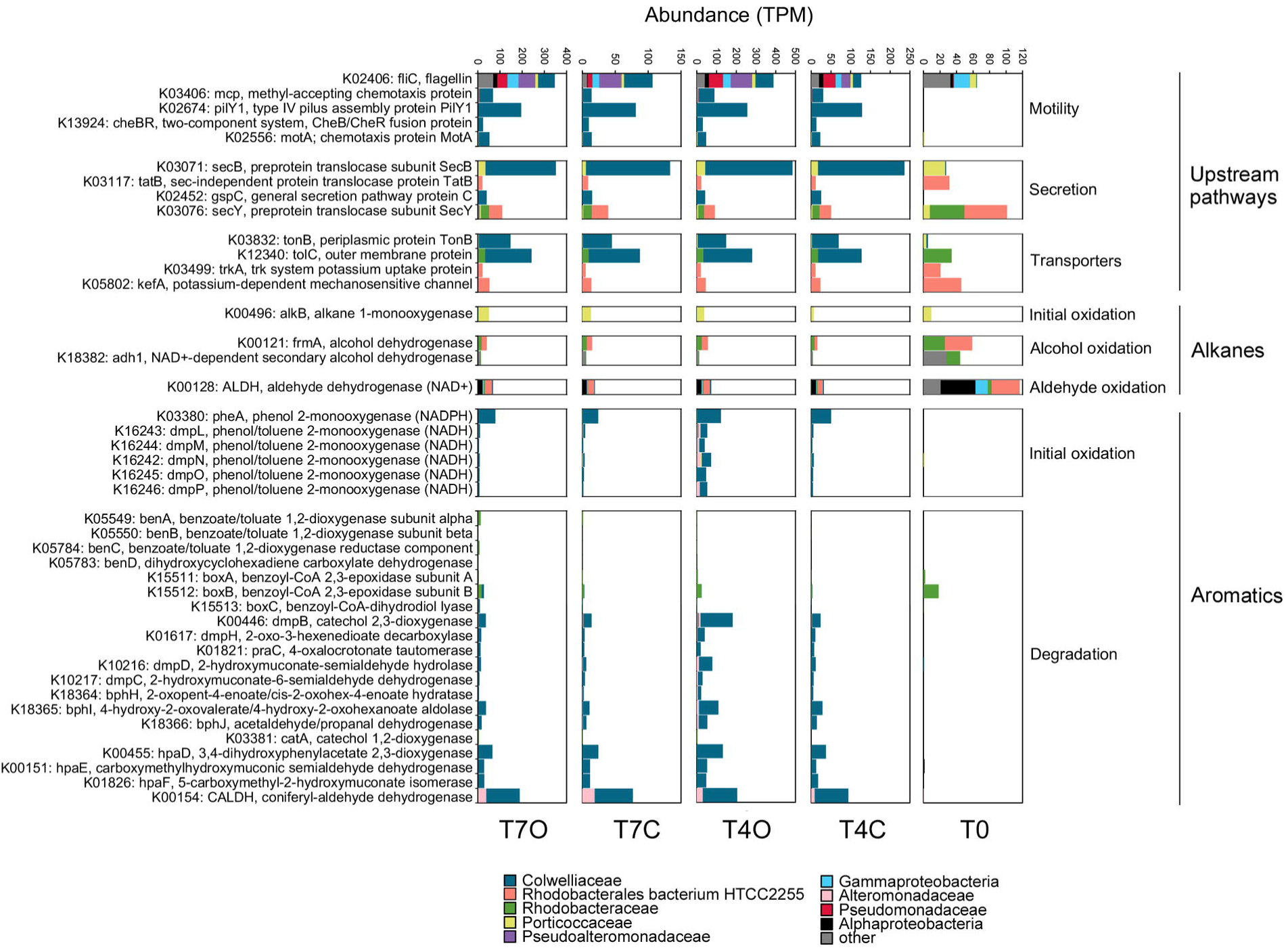
Taxonomic analysis of selected oil degrading genes. TPM values of upregulated genes (log2 fold-change ≥ 1, p ≤ 0.05) between control and oil exposed samples were grouped according to their gene function and colored by their taxonomic assignment.

#### 3.5.1 Upstream pathways: Motility, Secretion, and Transporters

The most abundant motility/flagella-related gene, *fliC*, was expressed mainly by the same bacterial families when comparing control and oil exposed seawater, however, the taxonomical profile of these 4 samples were clearly distinct from that of T0 (Figure 8). Notably, this gene was the most taxonomically diverse among the selected motility-related proteins. While T0 flagellins were dominated by Gammaproteobacteria, *Porticoccaceae*, Alphaproteobacteria and other unclassified bacteria, all other flagellins found in incubated samples were dominated by *Colwelliaceae, Pseudoalteromonadaceae* and *Pseudomonadaceae.* Flagellins with the highest log2-fold change (>5) and highest increase in TPM from day 4 to day 7 (T7O/T4O > 5) were from less abundant *Methylophaga* species, *Burkholderiaceae*, *Thiotrichaceae* and *Halomonadaceae* bacterium (Table S7). Other functions related to motility, e.g., *mcp*, *pilY*, *CheB/CheR* and *motA*, were all nearly absent from the T0 sample while being mostly expressed by the abundant *Colwelliaceae* family in all other samples. Among the selected secretion-related genes, *secB*, *tatB* and *gspC* exhibited the most dramatic shift in taxonomic composition with the increase in *Colwelliaceae* and *Alteromonadaceae* classified genes by day 4. All transporters annotated by KEGG showed a taxonomic shift from T0 to day 4 represented by the same two families, both in control and in oil exposed samples.

#### 3.5.2 Alkane degradation

The transcript abundance of alkane 1-monooxygenase (*alkB*) genes selected by the differential expression analysis was generally low (TPM sum < 40) and already detectable in the T0 sample. Out of the 11 DE genes annotated as *alkB* or *rubB*, eight were classified as SAR92 clade bacterium with an average log2-fold change of 2.33. This *Porticoccaceae* family bacterium contained both genes responsible for initial oxidation of n-alkanes (alkane 1-monooxygenase and rubredoxin NAD+ reductase) and was highly active in the acclimated seawater (T0) already. The highest log2-fold change was observed in case of a Gammaproteobacteria bacterium and a *Halieaceae* bacterium, but only the former showed an increasing trend from day 4 to day 7. However, no alkane monooxygenases from OHCB were found among the DE genes (Figure 9). Only 4 upregulated cytochrome P450 genes were found, with an average log2-fold change of 1.96 (classified as Bacteria and Gammaproteobacteria bacterium species).

**Figure 9.**
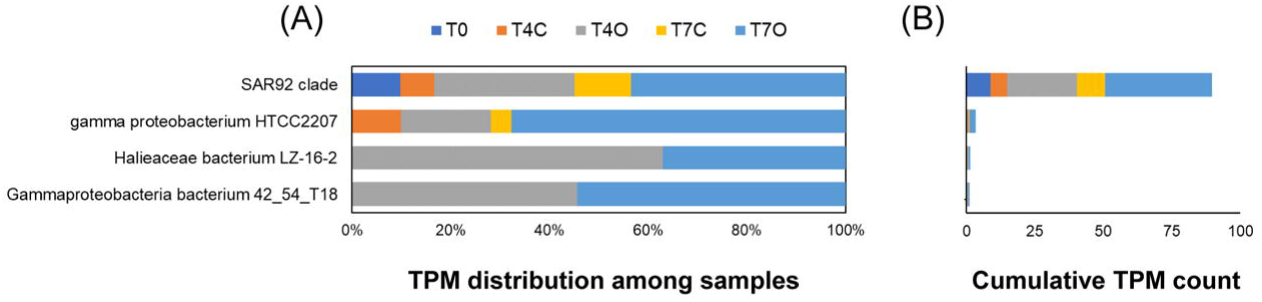
Upregulated alkane 1-monooxygenases. TPM distribution among the 5 samples (A) and cumulative TPM counts (B).

Many genes participating in the subsequent intermediary steps, *e.g.*, dehydrogenation of alcohols (90 genes), dehydrogenation of aldehydes (67 genes), and β-oxidation (134 genes, mainly from *Colwellia* sp.), were upregulated in oil exposed samples. Among these genes β-oxidation enzymes, *fadABJ* and ACSL were represented by the highest TPM counts, approximately 10-fold higher than the 11 oil-enriched alkane 1-monooxygenase genes. While the upregulated initial oxygenases were expressed almost exclusively by members of the family *Porticoccaceae* (mainly SAR92 clade bacterium) and unclassified Gammaproteobacteria, the β-oxidation genes were mainly transcribed by *Rhodobacteraceae* and *Colwelliaceae*. Abundant alcohol dehydrogenases *frmA*, *adh1* and also *qheDH* (last one not shown in Figure 8) showed taxonomic affiliation to Alphaproteobacteria (Rhodobacterales), *Pelagibacteraceae* and SAR92 clade, respectively. The highest TPM counts for these genes were observed in the T0 samples. The same was observed for the selected aldehyde dehydrogenase, ALDH, represented by taxa within Alphaproteobacteria. Both types of enzymes also maintained their taxonomic profile over time regardless of the presence of oil. Aldehyde dehydrogenases belonging to KO K00138 (*aldB*) however, were mostly absent from T0 (except for those associated with *Rhodobacterales* and *Pelagibacter*) and became enriched under laboratory conditions, in particular in the oil exposed tank (average log2-fold change = 2.2) (Table S7). These *aldB* genes were mainly associated with *Colwellia* species, *Arcobacter*, Rhodobacterales, Gammaproteobacteria and Candidatus *Pelagibacter*.

#### 3.5.3 Aromatic degradation

KEGG mapping of DE genes did not suggest the presence of aromatic ring-hydroxylating dioxygenases (RHDO) in the dataset reported here, however, BLAST annotated 29 DE genes as RHDO (Figure 10). The most abundant one was from *Rhodobacteraceae* bacterium and Rhodobacterales bacterium HTCC2255, both showing similar or somewhat higher TPM counts in oil exposed samples in comparison to the acclimated seawater (T0). Only a *Colwellia* sp. RHDO was oil-specific, however this gene had a very low TPM count.

**Figure 10.**
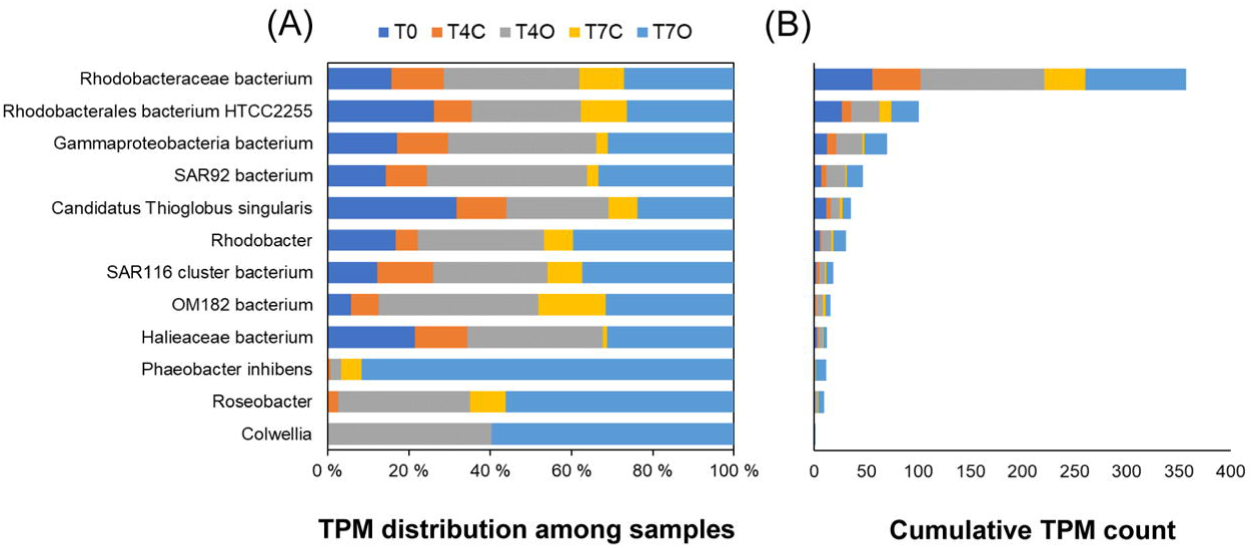
Upregulated ring-hydroxylating dioxygenases (RHDOs). TPM distribution among the 5 samples (A) and cumulative TPM counts (B).

KEGG analysis revealed an abundance of oxygenases and hydroxylases acting on monoaromatic compounds and highly expressed intermediary enzymes involved in channeling activated monoaromatic compounds towards the tricarboxylic acid (TCA) cycle. Enzymes responsible for initial oxidation, demethylation and subsequent conversion of monoaromatic rings were nearly absent from the T0 sample, except for *boxA* and *boxB* (two epoxidases involved in benzoate degradation, ring opening steps). Nearly complete pathways for several meta-cleavage routes of catechol degradation were reconstructed from the set of DE genes. Gene clusters for two different benzoate catabolic pathways were found: (1) ben pathway (*benABCD*) transcribed by *Rhodobacteraceae* and (2) box pathway (*boxABC*) transcribed mainly by *Rhodobacteraceae* and *Colwelliaceae*. Moreover, genes involved in aerobic benzene/toluene degradation (*dmpLMNOP*) transcribed mainly by *Colwelliaceae* were also enriched in the oil tank. Except for the *box* cluster, these aromatic compound converting enzymes were absent from the acclimated seawater but increased in the control tank as well. Moreover, by day 7, the number of transcripts responsible for aromatic compound degradation were reduced to levels similar to that in T4C. Contrary to other enzymes, benzoate/toluate 1,2-dioxygenase components and dihydroxy-cyclohexadiene carboxylate dehydrogenase (*benABCD*), and benzoyl-CoA converting enzymes (*boxABC*) maintained elevated TPM level in T7O. These enzymes cover three consecutive steps of the benzoate degradation pathway. Their taxonomic profiles were consistently dominated by the two most abundant families, *Colwelliaceae* and *Alteromonadaceae*.

To uncover highly oil-specific responses the DE genes were further explored. First, differentially abundant genes were ordered according to T7O/T4O ratio and only genes with a ratio > 10 were kept, corresponding to a ten-fold increase over time in oil exposed samples. Unannotated genes, hypothetical proteins, ribosomal proteins, and phage or virus transcripts were then removed leaving 294 genes for further evaluation. A large majority of the selected genes belonged to either *Methylophaga* species (96 genes) or Gammaproteobacteria bacterium (82 genes). The most abundant *Methylophaga* genes were 3-hexulose-6-phosphate synthase, F0F1 ATP synthase, elongation factor Tu, formaldehyde-activating enzyme and an ammonium transporter. For Gammaproteobacteria bacterium, a copper-binding protein, isocitrate lyase genes, F0F1 ATP synthase genes, formaldehyde-activating enzyme, benzoyl-CoA 2,3-epoxidase, transketolase genes, ammonium transporters, a porin, a cold-shock protein and a peroxiredoxin had the highest TPM in the T7O sample. Among the remaining genes, a phosphate starvation-inducible protein (*PhoH*), branched-chain amino acid ABC transporter substrate-binding protein of *Cycloclasticus*, together with TRAP transporter from *Colwellia*, ABC transporter and benzoate 1,2-dioxygenase from *Phaeobacter inhibens* had the highest TPMs (Table S7). Next, we focused on the OHCB associated DE genes and found the previously mentioned phosphate starvation-inducible protein (*PhoH*), a branched-chain amino acid ABC transporter substrate-binding protein of *Cycloclasticus*, ammonium transporters, flagellin, glutamine synthetase, lipoprotein NlpD, nitrate/nitrite transport system substrate-binding protein, nitrite reductase, nitrogen regulatory protein and urea transport protein of *Oleispira antarctica* and allophanate hydrolase from *Alcanivorax*. Finally, Venn diagram analysis of KEGG orthologs revealed that *tmoA* (toluene monooxygenase system protein), *benD* (dihydroxycyclohexadiene carboxylate dehydrogenase) and *kefB* (glutathione-regulated potassium-efflux system protein) were orthologues uniquely present in oil exposed seawater, albeit at very low TPM levels.

## 4 Discussion

### 4.1 Rapid decrease of petroleum hydrocarbon concentrations

Chemical analysis of seawater samples showed a rapid decrease in both aliphatic and aromatic hydrocarbon concentrations during the incubation. This likely resulted from a combination of abiotic and biotic processes, such as volatilization and coating to tank wall surface. Volatilization likely contributed to a significant portion of the loss of short-chain and monoaromatic hydrocarbons and naphthalenes. Adsorption to tank wall was observed. Compared to a study of Ribicic et al. (Ribicic, Netzer et al. 2018) the loss of PAHs occurred much earlier in our study (between day 1 and 3). They observed significant PAH biodegradation only between day 13 and 16 using North Sea, Norwegian fjord seawater, at a similar concentration of a chemically dispersed paraffinic oil (Statfjord crude, 3 mg·L^-1^). Possibly, a well-adapted microbial community could respond to oil exposure as quickly as suggested by the chemistry data in our experiment. Indeed, it has been shown that due to chronic influx of ubiquitous anthropogenic dissolved organic carbon (ADOC, including hydrocarbons) into the global ocean, natural bacterial communities evolved to quickly respond and taxa with recognized oil-degrading capabilities was shown to increase in relative abundance within 24 hr to ADOC amendment (Cerro-Gálvez, Casal et al. 2019). The microbial community of the seawater used in this experiment could have been partly adapted to low levels of oil contamination, although THC level was below detection limit in the acclimated seawater (T0). Nevertheless, three interesting signatures: (1) presence of active OHCB genera; mainly *Marinobacter* (0.17%), *Neptunomonas* (0.073%), *Oleispira* (0.060%), *Oleiphilus* (0.028%), *Thalassolituus* (0.0094%), and *Alcanivorax* (0.014%) (Figure 11), (2) expression of alkane monooxygenase and (3) ring hydroxylating dioxygenase (RHDO) genes in the T0 sample, may suggest some level of adaptation (Table S6).

**Figure 11.**
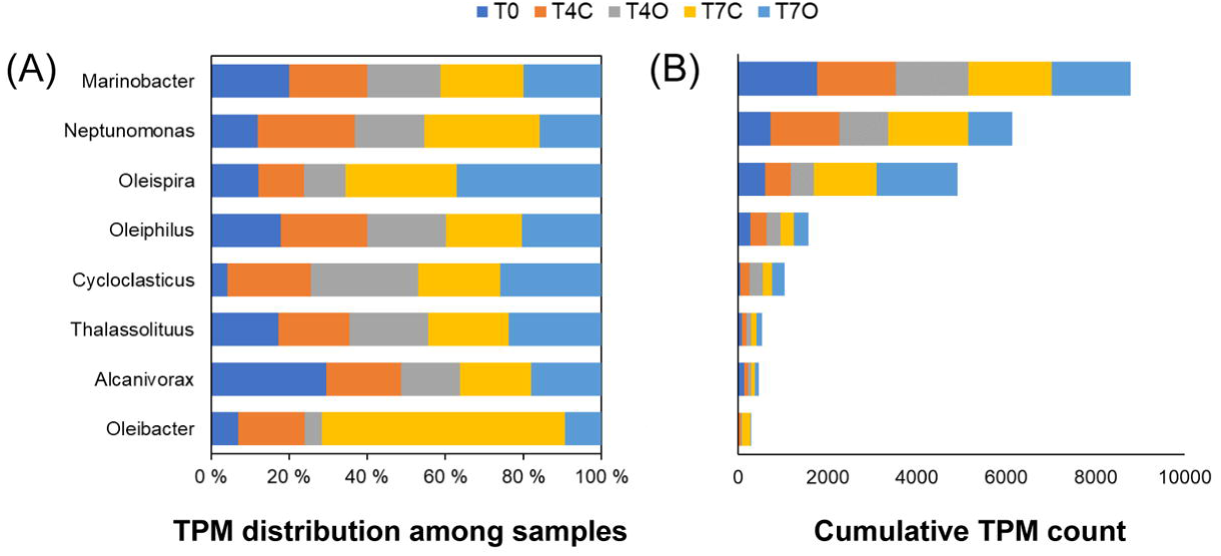
Obligate hydrocarbonoclastic bacteria (OHCB) detected in the metatranscriptome. Transcript per million (TPM) values (A) and distribution among the samples (B).

The very low relative abundance of OHCB genera, not only in the T0 sample but throughout the experiment, suggests that they did not represent significant biomass within the community, making it unlikely that they were responsible for rapid alkane and PAH biodegradation. Moreover, only *Oleispira* and *Alcanivorax*-associated alkane monooxygenases were present in the dataset, and only *Alcanivorax* (*Alcanivorax pacificus*) was expressing its alkane monooxygenase genes in the acclimated seawater sample, albeit at very low level (TPM of 1.021). Regarding ring-hydroxylating dioxygenases, none of these were classified into known OHCB genera. As Figure 11 shows, all the OHCB genera remained active or even became more active in the control tank compared to the oil exposed tank over time. Besides nearly identical activities in control and oil exposed seawater, their lack of increase in alkane monooxygenase relative expression in the presence of oil suggests that they were most likely utilizing carbon sources other than petroleum hydrocarbons (Radwan, Khanafer et al. 2019).

Other explanations for the rapid loss of both types of hydrocarbon compounds could include additional weathering processes such as absorption to biotic particles, e.g. phytoplankton. Phytoplankton were shown to be able to adsorb PAHs at high concentrations (up to 16 µg·g^-1^) into their biomass (Konat 1997) and phytoplankton absorption has recently been suggested as an important mechanism of PAH transport to sediments after their death (González-Gaya, Martínez-Varela et al. 2019). The metatranscriptomic analysis revealed that the acclimated seawater contained a large percentage of eukaryotic transcripts. As much as 8.4% of all BLASTX annotated predicted genes (48,273 out of 573,163) were of eukaryotic origin, with *Viridiplantae* being most dominant (61% of eukaryotic transcripts) followed by *Stramenopiles* (14%), *Opisthokonta* (10%), *Cryptophyta* (7%), *Haptophyceae* (5%), and *Alveolata* (2%). In comparison, 13% of transcripts were found to bin to similar eukaryotic groups during a phytoplankton bloom in the Gulf of Mexico (Rinta-Kanto, Sun et al. 2012). Although the absolute numbers of these organisms were not determined, their dominant presence in the T0 dataset suggests that influence of algal biomass should be taken into consideration when interpreting the chemistry results.

### 4.2 Microbial community and activities in the acclimated seawater

On the taxonomy side, the most abundant taxa in the acclimated seawater was the genus *Micromonas*, a green alga, while on the functional side, photosystem proteins from various eukaryotic phytoplankton, e.g. *Aureoumbra lagunensis*, *Teleaulax amphioxeia* and *Guillardia theta*, were the most relatively abundant proteins. Several bacterial taxa, often associated with phytoplankton-rich coastal marine communities, had comparable abundances to the dominant green algae and diatom groups in the acclimated seawater, *e.g. Polaribacter*, *Porticoccaceae* (possibly including SAR92), *Cryomorphaceae*, *Planktomarina*, *Formosa*, SAR86, *Crocinitomicaceae* and *Ulvibacter* (Table S5). These groups have been found during a diatom bloom in Monterey Bay, towards the end of a diatom-dominated spring phytoplankton bloom in the German Bight, in the bacterioplankton community of a *Phaeocystis globosa* spring bloom in the North Sea and in the bacterial community of a *Phaeocystis antarctica* bloom in the Amundsen Sea (Delmont, Hammar et al. 2014, Klindworth, Mann et al. 2014, Wemheuer, Wemheuer et al. 2015, Nowinski, Smith et al. 2019). Hence, the seawater collected for this experiment was indeed a typical example of a spring coastal marine microbial community. Interestingly, at least two of these genera, *i.e. Polaribacter* and *Ulvibacter*, are also considered to be typical examples of cold-adapted oil degrading bacteria (Brakstad, Lofthus et al. 2017).

It is well known that phytoplankton have a significant role in providing a habitat for hydrocarbon degrading bacteria (HCB) in the so-called photosphere (Suk and Mishamandani 2016, Thompson, Angelova et al. 2017). Phytoplankton can create hydrocarbon enriched zones in surface seawater as they can produce PAHs, volatile hydrocarbons, isoprene and hydrocarbon like substances such as alkenones (Kowalewska 1999, Binark, Güven et al. 2000). This phenomenon could in turn make phytoplankton rich seawater also more prepared to biodegrade accidentally spilled or leaked petrogenic hydrocarbons. While most green algae transcripts in the acclimated seawater belonged to typical coastal seawater green algae, e.g. *Micromonas* and *Bathycoccus*, we also found three genes of *Botryococcus braunii*, a well-known hydrocarbon-producing microalga (Jin, Dupré et al. 2016, Lopes dos Santos, Gourvil et al. 2016). This suggests the possibility that naturally occurring hydrocarbons could have been inducing the expression of the alkane monooxygenases observed in the T0 sample. *Crocinitomicaceae*, SAR92 clade and Bacteroidetes, which dominated the alkane monooxygenases expressed in the acclimated seawater (and other samples as well), and Rhodobacterales bacterium HTCC2255*, Halieaceae* and Candidatus *Pelagibacter ubique*, which dominated the RHDO transcripts in the acclimated seawater, have all been shown to be active in phytoplankton dominated communities and involved in the succession of phytoplankton blooms.

### 4.3 Dominant generalists and phytoplankton-associated taxa

Within three days of incubation, the entire microbial community changed dramatically both in the control and oil exposed tank, characterized by orders of magnitude decreases in algae and diatom transcripts concurrent with: (1) a significant decrease in mainly Bacteroidetes (e.g. *Flavobacteriaceae*, *Cryomorphaceae* and *Polaribacter*) and virus (e.g. *Phycodnaviridae*) transcripts, (2) a transient decrease in *Rhodobacteraceae*, and (3) a clear bloom of *Colwelliaceae*, *Campylobacterceae* (mainly *Arcobacter*) and *Alteromonadaceae*. The decline in algal and diatom transcript relative abundance could be associated with decreased activity due to insufficient light conditions or other limiting parameters in the incubation tanks. However, it is likely to have been a result of cell mortality due to viral infection. Taxonomic analysis of the viral contigs showed that excluding the unclassified viruses, the *Siphoviridae* and *Myoviridae* phage families dominated the viral community, followed by *Phycodnaviridae*, a family of viruses that infect eukaryotic algae (Table S9). *Phycodnaviridae* were particularly abundant in the acclimatized sample (T0), suggesting that they might have had a role in the decrease of *Micromonas* transcripts. If such a viral lysis event occurred, a significant amount of phytoplankton-derived organic material was released in both the control and oil exposed tanks, similarly to what could be observed at the end of a phytoplankton bloom – leading to the succession of opportunistic bacteria taking advantage of the abundant carbon source.

Indeed, one of the blooming bacterial taxa, *Colwellia,* has been previously associated with the degradation of phytoplankton-derived organic matter of a *Phaeocystis* bloom (Delmont, Hammar et al. 2014). *Colwellia* represents a group of heterotrophic cold-adapted bacteria able to degrade a wide range of organic compounds (Methe, Nelson et al. 2005). Through isolation from an enrichment culture, it has been demonstrated that some *Colwellia* have the capacity to degrade petroleum hydrocarbons in the presence of Corexit dispersant (Baelum, Borglin et al. 2012). The genome of this isolate has not been sequenced; therefore, the exact metabolic pathways are not known. Laboratory studies using stable isotope probing (SIP) showed that *Colwellia* can oxidize gaseous hydrocarbons (ethane and butane) (Redmond and Valentine 2012) and naphthalene (Sieradzki, Morando et al. 2019), but it is unclear whether they can utilize longer chain alkanes. *Colwellia* species have been suggested to be important players in the response to oil spills, however, it is well-known that there is a large variation among *Colwellia* species regarding their functional potential (Toseland, Moxon et al. 2014). This genus is often observed as the first responder in laboratory experiments, particularly under cold conditions (Redmond and Valentine 2012, Krolicka, Boccadoro et al. 2017). More commonly, however, *Colwellia* species co-occur with obligate hydrocarbonoclastic species, such as *Oleispira* or in later stages of oil degradation, with *Cycloclasticus* (Krolicka, Boccadoro et al. 2017, Ribicic, Netzer et al. 2018, Sieradzki, Morando et al. 2019). In this study, the nearly identical increase in the relative abundance of *Colwellia* transcripts in control and oil exposed seawater suggests that this bacterium, represented mainly by *Colwellia psychrerythraea* and Candidatus *Colwellia aromaticivorans*, utilized carbon from sources other than the added petroleum hydrocarbons such as carbon released from evanescent phytoplankton. Most recently, Candidatus *Colwellia aromaticivorans* was reported as a novel hydrocarbon-degrading bacterium identified through metagenomic analysis of a microcosm-scale oil exposure experiment (Campeao, Reis et al. 2017, Campeao, Swings et al. 2019). However, isolation and confirmation of this strain being able to grow on petroleum hydrocarbons has not been performed, only *in silico* predictions of its metabolic profile have been reported.

While *Campylobacteraceae* is also considered to be a group containing cold-adapted oil degrading bacteria, we observed a significantly lower relative abundance of *Campylobacteraceae* (in particular *Arcobacter*) in the oil exposed tank in comparison to the control tank. This might suggest that some *Campylobacteraceae* species were susceptible to toxic compounds present in crude oil rather than being involved in their degradation (Otth, Solís et al. 2005, Brakstad, Lofthus et al. 2017).

*Alteromonadaceae* represents a large group of motile Proteobacteria, which have been observed to form a transient bloom in response to manipulations during laboratory mesocosm set-up, where responses of bacterioplankton to either phosphorous or glucose addition were examined (Allers, Gómez-Consarnau et al. 2007). *Alteromonadaceae* also gathers a group of bacteria which have previously been reported as oil-degraders. On the one hand, the genus *Glaciecola* has been suggested as an early marker (responder) of oil pollution and an oil degrading *Glaciecola* strain has previously been isolated (Chronopoulou, Sanni et al. 2015, Krolicka, Boccadoro et al. 2017). On the other hand, this genus has also been suggested to be one of the dominant consumers of phytoplankton-derived organic matter during a cold-water spring bloom (von Scheibner, Sommer et al. 2017).

Surprisingly, none of the families with known OHCB genera became dominant during the incubation period of this study. In fact, our datasets showed an overall decline in abundance of the *Oceanospirillaceae* family, a group that has been often observed in oil spill response. However, one OHCB genus, *Oleispira* (*Oceanospirillaceae*), was slightly more abundant in oil-treated seawater by day 7. Krolicka et al. (Krolicka, Boccadoro et al. 2019) have suggested *Oleispira* to be a good bioindicator of oil occurrence in water based on the abundance and rapid response of this taxa shortly after oil contamination. The observed increase of *Oleispira* at day 7 in this study could suggest that the degradation of the added petroleum hydrocarbons was delayed until after the phytoplankton-derived organic matter was utilized by the opportunistic families. Alternatively, other non-OHCB taxa must have contributed to the initial alkane oxidation, such as SAR92 clade bacterium (*Porticoccaceae*). However, a previous study that analyzed metagenomes of microbial communities after exposure to oil showed that the family *Oceanospirillaceae* was contributing the most to initial *n*-alkanes degradation, whereas *Colwellia* and *Porticoccaceae* were involved in secondary *n*Lalkane catabolism and betaLoxidation (Ribicic, Netzer et al. 2018).

### 4.4 Dominant functions reflect the “tank effect”

Phosphorous acquisition dominated the microbial activity in the control tank (T4C and T7C samples), through uptake of dissolved inorganic phosphate and through transport of dissolved organic phosphate in the form of phosphonate. Phosphonate is an important component of the marine dissolved organic phosphorous pool and was suggested to represent a significant phosphorous source in the oceans (Villarreal-Chiu, Quinn et al. 2012). For the utilization of other forms of organic phosphorous, likely originating from dead phytoplankton, alkaline phosphatase was expressed (Shivaramu, Randive et al. 2019). This enzyme was found to be the most responsive process in a study where the effect of organic matter amendment was examined in a marine mesocosm (Baltar, Lundin et al. 2016). Dominance of phasin, a polyhydroxyalkanoate (PHA) granule-associated protein, possibly playing a role in PHA-related stress response, suggested the presence of excess organic matter which was likely utilized by bacteria for the synthesis of PHA for carbon and energy storage (Mezzina and Pettinari 2016). The high expression of the glyoxylate-shunt enzyme, isocitrate lyase, also suggests that organic carbon was rather utilized by incorporation into the biomass of rapidly growing opportunistic species (biosynthetic pathways) than by respiration to carbon-dioxide (Dunn, Ramírez-Trujillo et al. 2009). Overrepresentation of isocitrate lyase, together with the other glyoxylate shunt enzyme malate synthase, in a plume sample of the Deepwater Horizon spill was found by (Rivers, Sharma et al. 2013). According to the authors, this indicated the ongoing assimilation of hydrocarbon-derived intermediary metabolites (acetyl-CoA) into biomass.

The oil exposed tank (T4O and T7O samples) was characterized by the dominance of PQQ-dependent dehydrogenases (methanol/ethanol family) and aldehyde dehydrogenases (mainly from different *Colwellia* species). The TPM counts of these were at least an order of magnitude higher than initial oxygenases in the reported metatranscriptome (Table S6). An increase in PQQ-containing alcohol dehydrogenase expression was reported by (Rivers, Sharma et al. 2013) in samples retrieved from the plume of the DWH oil spill. They also reported higher expression of aldehyde dehydrogenases in the plume samples. However, in contrast with our results, this coincided with alkane monooxygenase expression and in general the levels of initial monooxygenase enzymes exceeded the levels of aldehyde dehydrogenases. Such co-occurrence was not observed in our dataset. Some PQQ-dependent ethanol dehydrogenases (*exaA*) and aldehyde dehydrogenases (*exaC*) taking part in the ethanol oxidation pathway, were shown to be essential for survival and growth at cold temperatures (Tribelli, Solar Venero et al. 2015). Hence, these dominant proteins could also reflect an essential aspect of the cold-adapted lifestyle of the active bacterioplankton in the oil exposed tank. Some TonB-dependent receptors (TBDRs), such as OmpS of *Alcanivorax dieselolei*, have been shown to be important in hydrocarbon transport across the bacterial cell envelop (Wang and Shao 2014). However, most of them are outer membrane proteins known for the active transport of iron siderophore complexes and acquisition of other large molecules, *e.g.* vitamin B12 (Blanvillain, Meyer et al. 2007). In addition, TBDRs could also be involved in transporting sugar oligomers (*i.e.* partly degraded polysaccharides) with the help of sugar-binding proteins (Matos, Lozada et al. 2016).

The major distinguishing characteristic of the T7O sample was the large amount of hypothetical proteins and phage/viral sequences. Although the total abundance of viral transcripts was similar over the course of the experiment, several phage-associated sequences showed a preferential increase in T7O. It is known that hydrocarbon pollution can increase virus abundance through prophage induction in marine bacteria (Jiang and Paul 1996, Cochran, Kellogg et al. 1998), leading to regulation of microbial population density by selective killing of abundant bacteria types (Thingstad 2000, Sauret, Bottjer et al. 2015), subsequently releasing back into the water dissolved organic carbon, which is then utilized by bacterial populations (Fuhrman 1992, Rosenberg, Bittan-Banin et al. 2010). Bacteriophages might also influence oil degradation by transferring hydrocarbon degradation genes between bacteria (Herrick, Stuart-Keil et al. 1997). Phage associated genes have been found to be more abundant in oil-contaminated seawater compared with uncontaminated water (Lu, Deng et al. 2012). A recent study showed that phages that infect known degraders can be highly abundant in hydrocarbon polluted water (Costeira, Doherty et al. 2019). Taxonomic analysis of the viral contigs at the species level showed abundance of the uncultured Mediterranean phage uvMED, abundant in the marine environment (Mizuno, Rodriguez-Valera et al. 2013, Garin-Fernandez, Pereira-Flores et al. 2018). Analysis of the contigs that showed > 2-fold increase in transcription in oil exposed samples, showed that most of the identified potential phage hosts belonged to *Pseudoalteromonas*, *Vibrio* and *Xanthomonas*. Their transcription increased from 4-8-fold on day 4 to 31-62-fold on day 7 (Table S9).

### 4.5 Upregulated functions

Oil pollution in the marine environment quickly triggers a number of microbial responses, including a broad range of activities: (1) chemotaxis, either away from toxic hydrocarbons or towards them, (2) efflux mechanisms to facilitate secretion of small hydrocarbons that may enter bacterial cells by passive transport, together with secretion systems to transport, proteins (toxins and extracellular enzymes) or biosurfactants outside the cell, (3) death and lysis of sensitive organisms which releases a pool of organic material available for the oil-tolerant and primary or secondary oil-degrading organisms, (4) adhesion to and biofilm formation on oil droplets and signaling between the biofilm forming cells, (5) an increase in nutrient transport and processes to generate available nitrogen, phosphorous and sulfur for growth on carbon-rich and nutrient-poor oil components, and (6) biosurfactant synthesis (Van Hamme, Singh et al. 2003, Mason, Hazen et al. 2012, Rivers, Sharma et al. 2013, Joye, Kleindienst et al. 2016, Parales and Ditty 2018). Indeed, most of these processes, also reported in recent large scale metatranscriptomic studies (Tremblay, Yergeau et al. 2017, Tremblay, Fortin et al. 2019), were represented in the upregulated genes in our dataset (Table S7). However, most of them were also increasing in expression levels in the control tank (Figure 8).

A straightforward explanation for the simultaneous increase in abundance in control and oil exposed tank could be that most of the upregulated genes were involved in the processing of phytoplankton-derived material released as a result of algal senescence during incubation. Many of the responses reported from bacteria in response to oil are similar to those reported in phytoplankton bloom succession studies and in experiments where anthropogenic dissolved organic carbon (ADOC) or deep-sea water (DSW) is added to seawater (Shi, McCarren et al. 2012, Klindworth, Mann et al. 2014, Cerro-Gálvez, Casal et al. 2019). Increased expression in response to these environmental perturbations typically includes the same broad categories of motility, chemotaxis, transporters, etc. Therefore, it can be challenging to identify truly specific indicators of anthropogenic oil contamination.

Functions associated with a declining phytoplankton bloom community were found in the presented dataset, including 283 differentially abundant genes encoding TRAP transporters (mainly from Alphaproteobacteria, *Rhodobacteraceae*, but also Candidatus *Colwellia aromaticivorans* and *Pelagibacteraceae*), 170 genes encoding branched-chain amino acid transporters (mainly from Alphaproteobacteria, *Rhodobacteraceae*, but also Gammaproteobacteria, *Colwellia*, and *Roseobacter*) and 55 genes encoding carbohydrate ABC-type transporters (mainly from Alphaproteobacteria, *Rhodobacteraceae* and *Oceanospirillaceae*). All these dissolved organic matter (DOM) uptake-associated functions were found to be significantly differentially represented in a bloom community and are strongly linked to the consumption of carboxylic acids, amino acids and carbohydrates released during bloom senescence (Rinta-Kanto, Sun et al. 2012). The presence of lysozyme genes from Candidatus *Colwellia aromaticivorans*, alginate lyases belonging to *Colwellia*, two glycoside hydrolases from *Glaciecola* and Alteromonadales and other enzymes, such methylamine dehydrogenases, taurine and sulfonate utilization genes among the upregulated genes further supports the hypothesis that some of the differentially abundant activities were likely reflecting response to phytoplankton-derived material (Suleiman, Zecher et al. 2016, Clifford, Varela et al. 2019, Landa, Burns et al. 2019).

Alkane 1-monooxygenases, rubredoxin, and P450 cytochromes have been suggested as important and specific functional markers of aliphatic hydrocarbon degradation (Wasmund, Burns et al. 2009, Scoma, Hernandez-Sanabria et al. 2015). However, the use of alkane 1-monooxygenase as marker for anthropogenic oil pollution in the marine environment remains controversial. Alkane 1-monooxygenase genes were found to be abundant in various seawater environments as determined by degenerate PCR (Wang, Wang et al. 2010) and metagenomic studies (Tremblay, Yergeau et al. 2017, Zhang, Xu et al. 2018, Tremblay, Fortin et al. 2019). One study that quantified *alkB* gene using qPCR found no clear distinction between the chronically oil polluted Baltic Sea coastal water compared to a nearby less exposed site (Miettinen, Bomberg et al. 2019). Similarly, metatranscriptomic analysis showed that alkane 1-monooxygenases had similar expression levels in plume samples in comparison to the non-plume following the Deepwater Horizon oil spill (Rivers, Sharma et al. 2013), while GeoChip-based functional gene array hybridization data showed significantly higher *alkB* gene numbers in the oil contaminated samples (19-26 genes) in comparison to non-contaminated ones (11-15 genes) (Lu, Deng et al. 2012). Yet another filed study found higher *alkB* gene expression, measured using qPCR, in oil-contaminated seawater of the Yellow sea in comparison to uncontaminated seawater (Wang, Zhang et al. 2014). In a laboratory experiment, Paisse et al (Paisse, Duran et al. 2011) observed that expression of *alkB* genes (reverse transcription-qPCR) was detected immediately after oil addition (after 1 hr) but diminished after 2 days, even though the alkane degradation was observed during the 14 days of incubation.

Aromatic ring-hydroxylating dioxygenases (RHDO, alpha and beta subunits) are used as robust markers of PAH biodegradation (González-Gaya, Martínez-Varela et al. 2019). Primers designed to amplify various RHDOs from Gram-negative and Gram-positive bacteria have been successfully used to differentiate oil-contaminated seawater of the Yellow sea from uncontaminated (Wang, Zhang et al. 2014). In our study, upregulated functions related to aromatic degradation were mostly associated with monooxygenases and dioxygenases involved in degradation of monoaromatic compounds (benzene, toluene, and phenol) and central intermediates (benzoate, catechol, protocatechuate, etc.). Monooxygenases and dioxygenases involved in aromatic degradation were also significantly enriched in the plume transcriptomes analyzed after the Deepwater Horizon oil spill (Rivers, Sharma et al. 2013). Among the enriched genes reported by Rivers et al., was gene cluster *benABC*, also found to be significantly upregulated on oil in our study (transcribed by *Rhodobacteraceae*). Most other upregulated aromatic monooxygenases (*dmpLMNOP* and *pheA*) and dioxygenases (*dmpB*, *catA* and *hpaD*) found in our study were transcribed by *Colwellia*, a genus that was abundant in both oil exposed and control incubations.

In order to establish how *alkB* genes or RHDOs could be used in a robust manner as oil pollution markers, further work is needed to consolidate the above-mentioned findings, particularly for *alkB*. It is important to keep in mind, that metatranscriptomics provides a taxonomic and functional profile of actively growing organisms and their expressed genes, resulting in knowledge about the relative contributions of active taxa and genes at a given time. However, an increase or a decrease in relative abundance of an organism or TPM of a gene does not necessarily mean that the absolute values of its biomass or its expression levels have changed. Quantitative approaches like qPCR are necessary to establish the absolute numbers of organisms or genes of interest. While metatranscriptomics is helpful in identifying gene targets, the robustness of these targets needs to be validated through quantitative assays and through field measurements thereafter. Further research needs to focus on linking together profiling-based approaches with quantitative ones and DNA-based methods with RNA-focused ones. In addition, quantitative analysis and profiling of dissolved organic matter in the seawater subjected to laboratory experiment or collected in the field, together with phytoplankton cell counts should be implemented in future studies in order to establish better control of these confounding factors.

## 5 Concluding remarks, implications and future work

This work was performed based on the assumption that the presence of oil in water can be rapidly evaluated using a set of functional genes expressed by microorganisms in response to oil. Such specific indicators of oil contamination could then be used on ecogenomic devices, such as the ESP, to detect oil contamination rapidly and monitor recovery in real time. Here, the metatranscriptome of the 5 samples archived by ESP instruments revealed some distinction between control and oil exposed samples, namely: (1) *Campylobacteraceae* (genus *Arcobacter*) was relatively less abundant in the oil treatment with the majority of the downregulated transcripts belonging to *Arcobacter* species, (2) a dominance (peak) of intermediary alcohol and aldehyde dehydrogenases, and monoaromatic ring-degrading enzymes on day 4 with a subsequent decline by day 7 except for benzoate/toluate 1,2-dioxygenase components and dihydroxy-cyclohexadiene carboxylate dehydrogenase (*benABCD*), and benzoyl-CoA converting enzymes (*boxABC*), and (3) a significant increase in phage-associated sequences, as well as *Methylophaga* and *Gammaproteobacteria* bacterium transcripts on day 7 in the oil treatment.

Furthermore, upregulated genes of OHCB, particularly *Cycloclasticus* (*i.e.* phosphate starvation inducible protein, branched-chain amino acid ABC transporter substrate-binding protein, ammonium transporter and glutamate synthetase), *Oleispira* (*i.e*. nitrite reductase, glutamine synthetase, urea transport and ammonium transporter) and *Alcanivorax* (allophanate hydrolase) were identified.

Both types of initial oxygenase enzymes, *e.g.* alkane monooxygenases and ring-hydroxylating dioxygenases responsible for the activation of aliphatic and aromatic hydrocarbons, were present already in the acclimated seawater. Only 11 out of 86 alkane monooxygenases and 29 out of 427 aromatic ring-hydroxylating dioxygenases were upregulated in oil exposed samples in comparison to control. Initial monooxygenases from OHCB were found, however, only for alkane monooxygenase (*Oleispira* and *Alcanivorax*) and not among the upregulated genes.

Despite the observed differences, we conclude that the effect of the relatively low concentration oil exposure became largely masked and/or perhaps delayed, likely due to the simultaneous release and degradation of algae-derived organic matter. Abundant genes, *e.g.* alcohol and aldehyde dehydrogenases, which differentiated oil exposed seawater from control, were likely involved in processing of phytoplankton-derived material. Our results therefore highlight the importance of considering the overall taxonomic context and interactions within all domains of life present in an ecosystem, when interpreting the effect of oil exposures on bacterial communities. Moreover, experimental designs for discovering and testing functional markers need to incorporate appropriate controls not only for potential abiotic confounding factors (temperature, light, nutrients, *etc*.) but also biotic ones, *e.g.*, the response of microeukaryotes and viruses to laboratory enclosure. This study emphasizes that careful considerations in experimental design are needed when studying the effects of contaminants on community level functional profiles, and that there is a need to validate laboratory findings with actual field conditions. Furthermore, given the complexity of the interactions between ecosystem members, network analysis could uncover an array of functional genes, which together implemented as a multitarget assay, could be used in sensitive and robust warning systems.

Further insights into taxonomic and functional changes of microbial communities inferred from experimental work that consider environmental factors such as seasonality and organic matter content are needed. Thereafter, a robust array of targets best appropriate for integration into genosensor technologies, such as ESP, can be selected and used for sensing community changes in response to oil contamination.

## 6 Data Availability

The datasets generated for this study can be found in the NCBI’s Sequence Read Archive (SRA). The raw sequencing reads generated through amplicon sequencing are publicly available under the project number PRJEB32487 (ERS3402758-ERS3402761). The raw sequencing reads generated through metatranscriptomics are publicly available under the project number PRJNA556155.

## 8 Author Contributions

TB designed and supervised the study. TB and AK performed the mesocosm experiment and sampling. KK extracted DNA and RNA and arranged the RNA sequencing. AK arranged the amplicon sequencing and performed the analysis of the amplicon sequencing data. KK and AB performed the analysis of the metatranscriptomic data. AB, KK, and TB prepared the manuscript. AK revised the manuscript. KK and AB contributed equally to this work.

## 10 Acknowledgments

We are very thankful to the MBARI ESP team and particularly Dr. Christina Preston and Roman Marin III for their scientific expertise and technical contribution during ESP work. We wish to dedicate this paper to Roman Marin III who recently passed away. We also would like to thank our colleagues at NORCE, Mark Berry, Stig Westerlund, Sreerekha S. Ramanand and Kjell Birger Øysæd for their contribution to the experimental work and as well Emily Lyng for manuscript proof-reading.

