## Supplementary material for "Discovery of functional gene markers of bacteria for monitoring hydrocarbon pollution in the marine environment - a metatranscriptomics approach": Table S1

**Table S1.** Sample filtration volume and RNA quantification of samples sequenced in this study.

|  | T0 | T4C | T4O | T7C | T7O |
| --- | --- | --- | --- | --- | --- |
| Sample filtration volume (ml) | 1000 | 700 | 700 | 700 | 700 |
| RNA concentration (ng/µl) | 57 | 216 | 201 | 78 | 177 |
| Total RNA sent (µg) | 2.3 | 8.6 | 8.0 | 3.1 | 7.0 |
