## Supplementary material for "Discovery of functional gene markers of bacteria for monitoring hydrocarbon pollution in the marine environment - a metatranscriptomics approach": Table S2

**Table S2.** Statistics of sequence processing.

|  | T0 | T4C | T4O | T7C | T7O |
| --- | --- | --- | --- | --- | --- |
| Raw reads (Million) | 37.1 | 52.5 | 54.9 | 49.9 | 48.7 |
| Total bases (Gb) | 2.8 | 3.96 | 4.15 | 3.76 | 3.67 |
| Read length (bp) | 75 | 75 | 75 | 75 | 75 |
| Quality filtered reads | 17,816,080 | 28,530,130 | 29,020,210 | 27,545,407 | 23,999,441 |
| Number of assembled contigs | 628,164 | | | | |
| Number of predicted CDS | 674,434 | | | | |
| Reads mapped back to contigs | 6,185,558 | 19,813,474 | 19,903,786 | 18,641,091 | 17,069,785 |
| Reads mapped back to CDS | 4,738,252 | 16,890,778 | 16,906,056 | 15,901,564 | 14,358,262 |
