## Supplementary material for "Discovery of functional gene markers of bacteria for monitoring hydrocarbon pollution in the marine environment - a metatranscriptomics approach": Table S4

**Table S4.** Diversity metrics calculated on family level from the RNA sequencing (RNAseq) and amplicon sequencing (16S) datasets. H – Shannon index and J – Pielou’s evenness.

|  | T0 | | T4C | | T4O | | T7C | | T7O | |
| --- | --- | --- | --- | --- | --- | --- | --- | --- | --- | --- |
|  | RNAseq | 16S | RNAseq | 16S | RNAseq | 16S | RNAseq | 16S | RNAseq | 16S |
| *H* | 2.108 | n.a. | 1.935 | 1.971 | 1.978 | 1.740 | 1.973 | 1.934 | 1.979 | 1.714 |
| *J* | 0.418 | n.a. | 0.401 | 0.484 | 0.406 | 0.430 | 0.405 | 0.485 | 0.409 | 0.420 |
