## Supplementary figures and images for "Discovery of functional gene markers of bacteria for monitoring hydrocarbon pollution in the marine environment - a metatranscriptomics approach"

### Figure S1

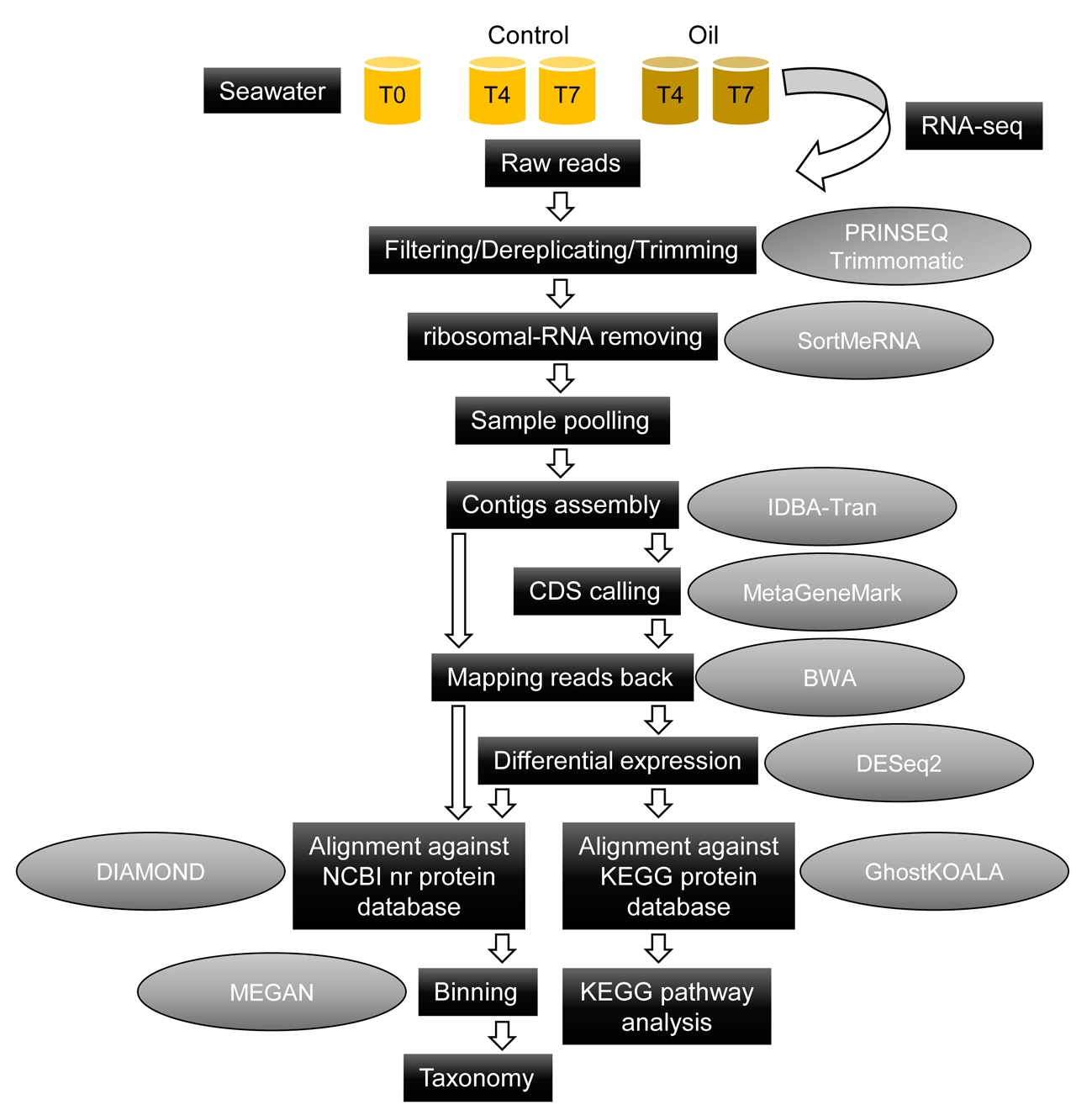
